# enmpa: An R package for ecological niche modeling using presence-absence data and generalized linear models

**DOI:** 10.1101/2024.02.20.581032

**Authors:** Luis F. Arias-Giraldo, Marlon E. Cobos

**Author notes:** The authors contributed equally to this work. Corresponding author: Luis F. Arias-Giraldo.

## Abstract

Here, we present the new R package “enmpa,” which includes a range of tools for modeling ecological niches using presence-absence data via logistic generalized linear models. The package allows users to calibrate, select, project, and evaluate models using independent data. We have emphasized a comprehensive search for ideal predictor combinations, including linear, quadratic, and two-way interaction responses, to provide more detailed and robust model calibration processes. We demonstrate the use of the package with an example of a simulated pathogen and its niche. Since enmpa is designed specifically to work with presence-absence data, our tools are particularly useful for studies with data derived from a detection or non-detection sampling universe, such as pathogen testing results. enmpa can be downloaded from CRAN, and the source code is freely available on GitHub.

## Introduction

Ecological niche modeling (ENM), often also referred to as species distribution modeling (SDM), constitutes a range of analytical methods employed extensively in ecological research (Guisan and Zimmermann 2000; Franklin 2010; Peterson et al. 2011). These methods have been proven particularly useful in characterizing and predicting Grinnellian (abiotic or non-interactive) ecological niches of species. Applications of these methods span various fields, including conservation planning (Franklin 2013; Hannah et al. 2020), climate change impact assessment (Searcy and Shaffer 2016; Blowes et al. 2019), potential biological invasions (Jiménez-Valverde et al. 2011; Park and Potter 2015; Cordier et al. 2020), and disease risk mapping (Peterson 2014).

Several modeling methods are available within the ENM framework, which can be classified by the types of data that they use: presence-only, presence and background (or “pseudoabsence”), and presence-absence data (Elith et al. 2006; Peterson et al. 2011). More generally, methods for ENM fall into three broad categories: ‘profile,’ ‘regression,’ or ‘machine-learning.’ Profile methods consider presence data only. Regression and machine-learning methods use both presence-absence or presence-background data.

An essential consideration in ecological studies is understanding the meaning of the outputs generated by algorithms. A common objective of these studies is to model the probability of presence of a particular species of interest. However, it is crucial to note that estimating probabilities of occurrence requires rigorous comparisons of presence and absence data (Ward et al. 2009). Modeling applications that utilize presence-only data can, at best, estimate relative suitability (Ferrier et al. 2002). Although availability of occurrence data poses a significant challenge in ecological niche modeling, biodiversity data-portals like the Global Biodiversity Information Facility (GBIF) can serve as valuable sources of occurrence data records, at least potentially including presences and absences.

Among the different modeling methods that are used in ENM, generalized linear models (GLMs), an extension of classical multiple regression, have yield reliable results in ecological research in estimating probability of occurrence of species (Guisan et al. 2002; Bolker et al. 2009; Rupprecht et al. 2011; Ghanbarian et al. 2019). Determination of species’ responses to environmental gradients is of particular interest for most biological questions in ENM, which is given by the shapes of response curves (Guisan et al. 2002; Oksanen and Minchin 2002; Austin 2007; Santika and Hutchinson 2009). Fitting GLMs via a logistic link function for presence-absence binomial data offers results approximating Gaussian curves according to the principles of ecological niche theory (Austin 2007; Santika and Hutchinson 2009). However, response curves in GLMs may be inappropriate or unrealistic when models are not tuned adequately (Austin et al. 1990).

It is crucial to explore and determine appropriate parameter settings for models, instead of simply using default settings (Warren and Seifert 2011). This task can be accomplished through model calibration and selection processes (Radosavljevic and Anderson 2014; Hao et al. 2020). Models resulting from calibration exercises help to describe the phenomenon of interest better while achieving a robust fit to the data, high predictive performance, and generalizable model terms (Cobos et al. 2019a). To our knowledge, well-defined tuning routines have yet to be developed for GLMs in ENM, unlike methods like Maxent (Phillips et al. 2006; Muscarella et al. 2014; Phillips et al. 2017; Cobos et al. 2019a).

To bridge this methodological gap (GLM tuning routines for ENM), we introduce the enmpa R package (R Core Team 2022). This package is designed to refine the calibration and parameter tuning processes, offering a solution to the challenges in modeling robust ecological niches with presence-absence data. Via enmpa, we explore the entire range of possible model configurations to identify the most suitable and practical parameterizations for ENMs.

### Package Description

The enmpa R package provides a set of tools to automate various ENM steps using logistic regressions via GLMs, focusing on fitting linear and quadratic relationships and multiplicative interactions between predictors. The response (dependent) variable is a set of presence and absence records, and the predictor (independent) variables can be defined according to the question (e.g., bioclimatic variables). Major steps enabled via enmpa include model calibration (candidate model fitting and evaluation), model selection, model transfers, and model evaluation with independent data.

### Data required

The input data required consist of presence-absence records associated with values of the independent variables. The dependent variable is the set of presence-absence observations. The data must be structured as a data.frame in R to ensure the proper functioning of the package.

### Exploration of variables for models

We adapted methods developed by Cobos and Peterson (2022) to identify relevant variables for characterizing species’ ecological niches. These methods include two complementary statistical analyses: (1) a multivariate approach based on a permutational multivariate analysis of variance (PERMANOVA) (Anderson 2017) and (2) a univariate non-parametric method based on descriptive statistics and randomizations. These methods are designed to allow characterizations of signals of ecological niches while considering the sampling universe explicitly. As enmpa uses presence-absence data through logistic regression, we consider it appropriate to implement this method as a potential variable selection step prior to modeling. Together with proper considerations of variable biological relevance, this step can help to reduce initial numbers of predictors considered for ENM analysis (Cobos et al. 2019b).

### Model calibration

The model calibration step aims to determine which combination of parameter settings best represents the phenomenon of interest via exploration of performance metrics that characterize how well models fit the data (Steele and Werndl 2013). The tools in enmpa automate a process that includes three main steps: (1) fitting candidate models with distinct parameter settings, (2) evaluating their performance, and (3) selecting the most robust candidates based on predefined criteria.

#### Candidate model fitting

In this package, we propose exploring distinct parameter settings by producing multiple model formulas to fit models to the data. These formulas derive from combinations of predictors that can be obtained using the original independent variables and response types: linear (l), quadratic (q), and product (p). This approach allows exhaustive exploration of all possible combinations of predictors, enabling a detailed analysis of the entire predictor setting space (Cobos et al. 2019b). Users can produce all these formulas manually or use functions in enmpa designed to automate the process considering two main inputs: variable names and response types. Users can also define the permutation strategy to create formulas according to the desired level of intensiveness in exploring setting options (e.g., only increasing complexity of variable combinations, or all independent and combinatorial options).

#### Candidate model evaluation

To evaluate candidate models, we use three complementary approaches. The first tests predictive power using a *k*-fold cross-validation approach (Hastie et al. 2009). The original dataset is partitioned into *k* subsets (folds) aiming for equal size and maintaining the original prevalence (ratio of presences and absences). The algorithm performs *k* iterations of training, in which each iteration uses *k* - 1 folds for training and keeps one fold for testing. This process evaluates model discrimination and classification capacities.

Discrimination is measured using the area under the receiver operating curve (ROC-AUC), a non-threshold dependent metric used, in our case, solely to detect models that perform better than random expectations (Lobo et al. 2008). Classification ability is measured via several metrics deriving from an estimated confusion matrix, including sensitivity, specificity, accuracy, false positive rate, and true skill statistic (TSS), all of them according to three thresholds (i.e., equal sensitivity and specificity, sensitivity of 90%, and maximum TSS) (Fielding and Bell 1997; Manel et al. 2001; Allouche et al. 2006; Liu et al. 2011). Means and standard deviations of these metrics are calculated to summarize the model’s predictive performance and help to select the best models.

The second approach uses the Akaike Information Criterion (AIC) (Akaike 1998; Warren and Seifert 2011; Warren et al. 2014) to assess model goodness-of-fit, accounting for model complexity. This metric offers a relative quality measure to other candidate models based on the same dataset with different parameters. AIC increases with information loss, so the best model for a set of occurrence data is the one with the lowest AIC. To compare AIC values of multiple models directly, we also calculate ΔAIC (Wagenmakers and Farrell 2004) by subtracting the AIC of the best model (the one with the lowest AIC) from the AIC of each model being compared, as follows: ΔAIC_i_ = AIC_i_ − AIC_min_. A ΔAIC of 0 indicates that the model in question is the best model; models with ΔAIC values ≤2 have substantial support, and can be considered almost as good as those with the lowest AIC. Subsequently, Akaike weights are derived from ΔAIC for model averaging, representing a model’s relative likelihood: Akaike weight (W_i_) is calculated as an exponent of the negative half of its ΔAIC value, and the relative likelihoods are normalized so that their sum across all models compared equals 1. This later step is achieved by dividing the relative likelihood of each model by the sum of the relative likelihoods for all models, producing the Akaike weight for each model. We note that the implementation of AIC calculations for GLM consider models goodness-of-fit, whereas that for maxent, which has seen considerable use, AIC is based on the model predictions (Warren and Seifert 2011).

Finally, enmpa incorporates an extra evaluation step involving analyzing the response curves of quadratic terms. Quadratic features are optimal in studies of responses of species to variable gradients, and fit well with ecological niche theory (Austin 2007; Santika and Hutchinson 2009). However, a limitation of quadratic features is that they can be concave upward, yielding a binomial response, which does not fit well with theory (Austin 2007). By investigating the coefficients of a second-degree equation (*y = β_0_ + β_1_x + β_2_x^2^*), we can infer the shape of the curve. A positive *β_2_* suggests a *U*-shaped bimodal curve, while a negative *β_2_* suggests a Gaussian-shaped unimodal curve. As such, we implemented a filter to retain only those candidate models in which any quadratic responses are *β_2_*-negative.

#### Model selection

To choose the best candidate models, we follow a set of criteria, which are prioritized as follows: (1) we only consider models with ROC-AUC > 0.5; (2) from among such models, we only keep those that have an acceptable predictive ability (TSS ≥ 0.4); and (3) from among all the models passing the first two filters, we chose those with good fitting and appropriate complexity, those with ΔAIC ≤ 2 (Burnham and Anderson 1998).

In addition, enmpa includes an optional filter that takes into account the shape of quadratic response curves. Users have the option to consider only models with predictors that have unimodal responses. If this filter is used, it is applied before the first three filters. Even if the bimodal response of species could have interesting and meaningful interpretations, it is tricky to determine optima and understand species’ tolerances, which need to be evaluated individually for particular species. Therefore, we recommend considering only models with unimodal or monotonic responses.

### Variable contribution and response curves

Variable contribution and analysis response curves are model outputs with practical relevance to researchers interested in interpreting model outputs. Two well-established methods for determining the importance of predictor variables in GLMs are used to evaluate the individual contributions of variables (Murray and Conner 2009) and visualize predicted responses of species to specific predictor variables (Elith et al. 2005).

A response curve represents the relationship between the probability of occurrence of a species (dependent variable) and environmental variables (independent variables) included in the model. This curve describes the predicted probabilities across a range of values for a given environmental variable. The enmpa package calculates the probabilities along a single environmental gradient to estimate response curves while holding all other gradients constant at their mean values (Elith et al. 2005).

To identify the most significant predictors in our models, we use a variable contribution analysis based on the deviance explained by predictors relative to the complete model deviance (Guisan and Zimmermann 2000; Clouvel et al. 2023). Deviance is a measure of the model’s lack of fit, so a decrease in deviance indicates an improvement in model fit when a predictor variable is included. To assess predictor importance, then, we implemented the following procedure: (1) a GLM including all predictors is fitted; (2) the initial deviance is calculated; (3) each predictor is removed iteratively from the model, and a new deviance is calculated; (4) the decrease in deviance after adding each predictor is calculated; (5) the decrease in deviance is normalized and expressed as a ratio; and (6) predictors are ranked based on how they help to decrease model deviance: variables with higher ratios are considered more important for the model.

### Model projections

The enmpa package facilitates transferring selected models to different areas or scenarios, with two options: free extrapolation and extrapolation with clamping. These options are available for all variables or for just selected variables. When using free extrapolation, predictions will follow the response patterns when variable values are outside the ranges of the environmental data under which models were calibrated. In contrast, extrapolation with clamping limits the response to the level manifested at the boundaries of calibration values.

### Model consensus

One way of selecting a model is to choose the “best” one for the data based on one or a set of predictive performance metrics (Elith et al. 2006). However, an alternative method is to use a consensus of models (Thuiller 2003; Qiao et al. 2015). Consensus approaches may provide more robust predictions by leveraging the general agreement among models with similar performance. A consensus result can be calculated as the mean, median, or weighted average of any set of models. Mean and median results are calculated using the predictions of all selected models, whereas a weighted average is calculated using these predictions and the AIC weights for each model. Models with higher AIC weights contribute more to the consensus when the weighted average option is used.

### Model evaluation with independent data

Ideally, the final model should be evaluated using independent test data, although it is often challenging to find data independent from those used to create the models (Araújo and Guisan 2006; Peterson et al. 2011). In cases in which independent data are available, enmpa facilitates evaluation of final model predictions. Users can provide presence-absence or presence-only records for this evaluation. Evaluation in cases involving presence-only data includes partial ROC and omission error (*E*) metrics (Cobos et al. 2019a). For presence-absence data, the metrics returned are the ROC AUC, sensitivity, specificity, accuracy, false positive rate, and true skill statistic (TSS), and for three thresholds (maximum TSS, equal sensitivity and selectivity, and a value that ensures a sensitivity of 90%).

### Example application

Here, we provide a guide on how to use enmpa, in the form of a worked example. This example includes the processes of variable exploration, model calibration and selection, transfers to specific areas and scenarios of interest, and post-modeling analyses. The code required to reproduce this example can be found in a GitHub repository (available at https://github.com/Luisagi/enmpa_test). The occurrence data, raster layers, and an independent test dataset used in these examples are included in the package (available at https://CRAN.R-project.org/package=enmpa).

### Data

The example data include 500 presence records of a virtual pathogen detected in a total of 5627 virtual host samples (i.e., 5127 absences), for a prevalence of 8.9%. The example data was generated based on ellipsoidal virtual niches for the host and the pathogen using the package evniche (https://github.com/marlonecobos/evniche) in R v4.2.2 (R Core Team 2022). The pathogen was designed to have a higher prevalence towards warm and dry environmental conditions. Example data to illustrate independent tests were generated the same way but during a posterior set of analysis. The environmental data related to occurrence records were extracted from raster layers that represent two Bioclimatic variables (annual mean temperature and annual precipitation, BIO-1 and BIO-12), from the WorldClim database v2.0 (Fick and Hijmans 2017). The two datasets associated with environmental values are included as part of example data in enmpa.

### Analysis

#### Variable exploration for niche signal detection

We started with exploratory analyses to detect whether the environmental variables can describe the niche of the virtual pathogen (Cobos and Peterson 2022). We conducted the two tests (multivariate and univariate approaches) to assess whether the position and spread of the pathogen niche differed from those of the host, considering the two environmental variables in the example.

#### Calibration and model selection

In all 31 candidate models were created using combinations of the two environmental variables, with linear, quadratic, and product responses. Each model was tested for model complexity (AIC) using the whole dataset and with a *k*-fold cross-validation (*k* = 5) to evaluate its performance in terms of discrimination (ROC-AUC) and classification capacity (false positive rate, accuracy, sensitivity, specificity, and TSS). The classification metrics were based on three relevant thresholds: equal sensitivity and specificity (ESS), the sensitivity of 90% (SEN90), and maximum TSS (maxTSS). The best models were selected according to enmpa criteria: only models with only convex quadratic responses were assessed, then, models with ROC-AUC > 0.5 were retained, and after that, only models with a good classification capacity (TSS > 0.4) were selected. Finally, only the models with ΔAICc values ≤ 2 were chosen as the final selected models.

#### Post-modeling analyses

We analyzed the contributions of predictors to the model and explored variable response curves for each model and the consensus results. Projections to the 48 contiguous United States were produced for all models selected using the aforementioned criteria. We used three approaches to generate consensus results: mean, median, and weighted average. To represent model variability, we also calculated the variance among selected models. Finally, the independent data were used to evaluate the models selected and the consensus results.

### Example results

Both tests for environmental sensitivity consistently performed well for the two bioclimatic variables for the virtual pathogen case. Multivariate analysis based on PERMANOVA effectively detected niche dissimilarities between the pathogen and host, with the pathogen niche forming a subgroup nested within the host niche (Fig. S1). Using the mean as the comparison metric for the univariate non-parametric test, we found that the pathogen niche was shifted towards high temperature and lower precipitation values. Comparisons using the standard deviation showed that the pathogen niche was narrower than the host niche in both variables (Fig. S2).

After model calibration, eight models were excluded because they presented concave quadratic responses. The remaining 23 models met the next criterion with a ROC-AUC larger than 0.5 (Table 2). Of these models, 19 performed well, with a TSS larger than 0.4. Finally, only two models met all of the evaluation criteria regarding discrimination, prediction ability, and fitting considering complexity (ΔAIC ≤ 2).

**Table 1.**
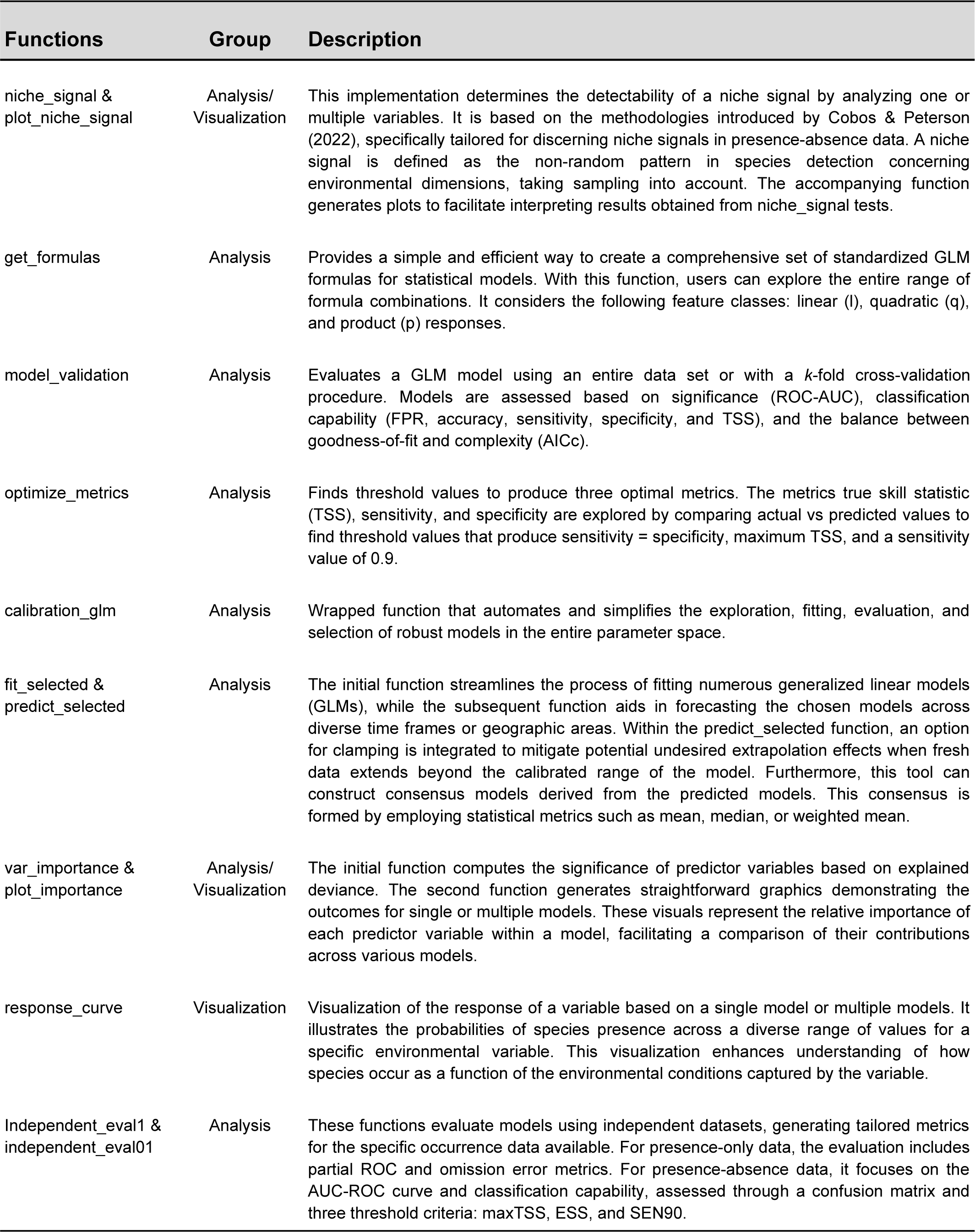
Description of the main functions included in the R package enmpa.

| Functions | Group | Description |
| --- | --- | --- |
| niche_signal & plot_niche_signal | Analysis/<br>Visualization | This implementation determines the detectability of a niche signal by analyzing one or multiple variables. It is based on the methodologies introduced by Cobos & Peterson (2022), specifically tailored for discerning niche signals in presence-absence data. A niche signal is defined as the non-random pattern in species detection concerning environmental dimensions, taking sampling into account. The accompanying function generates plots to facilitate interpreting results obtained from niche_signal tests. |
| get_formulas | Analysis | Provides a simple and efficient way to create a comprehensive set of standardized GLM formulas for statistical models. With this function, users can explore the entire range of formula combinations. It considers the following feature classes: linear (l), quadratic (q), and product (p) responses. |
| model_validation | Analysis | Evaluates a GLM model using an entire data set or with a *k*-fold cross-validation procedure. Models are assessed based on significance (ROC-AUC), classification capability (FPR, accuracy, sensitivity, specificity, and TSS), and the balance between goodness-of-fit and complexity (AICc). |
| optimize_metrics | Analysis | Finds threshold values to produce three optimal metrics. The metrics true skill statistic (TSS), sensitivity, and specificity are explored by comparing actual vs predicted values to find threshold values that produce sensitivity = specificity, maximum TSS, and a sensitivity value of 0.9. |
| calibration_glm | Analysis | Wrapped function that automates and simplifies the exploration, fitting, evaluation, and selection of robust models in the entire parameter space. |
| fit_selected & predict_selected | Analysis | The initial function streamlines the process of fitting numerous generalized linear models (GLMs), while the subsequent function aids in forecasting the chosen models across diverse time frames or geographic areas. Within the predict_selected function, an option for clamping is integrated to mitigate potential undesired extrapolation effects when fresh data extends beyond the calibrated range of the model. Furthermore, this tool can construct consensus models derived from the predicted models. This consensus is formed by employing statistical metrics such as mean, median, or weighted mean. |
| var_importance & plot_importance | Analysis/<br>Visualization | The initial function computes the significance of predictor variables based on explained deviance. The second function generates straightforward graphics demonstrating the outcomes for single or multiple models. These visuals represent the relative importance of each predictor variable within a model, facilitating a comparison of their contributions across various models. |
| response_curve | Visualization | Visualization of the response of a variable based on a single model or multiple models. It illustrates the probabilities of species presence across a diverse range of values for a specific environmental variable. This visualization enhances understanding of how species occur as a function of the environmental conditions captured by the variable. |
| Independent_eval1 & independent_eval01 | Analysis | These functions evaluate models using independent datasets, generating tailored metrics for the specific occurrence data available. For presence-only data, the evaluation includes partial ROC and omission error metrics. For presence-absence data, it focuses on the AUC-ROC curve and classification capability, assessed through a confusion matrix and three threshold criteria: maxTSS, ESS, and SEN90. |

**Table 2.** Model calibration summary. The table displays the evaluation of the main metrics for the 31 candidates evaluated using a cross-validated *k*-fold (*k* = 5) analysis. The most robust models selected using three criteria implemented in enmpa are indicated in bold. ROC-AUC, sensitivity, specificity, and TSS values are presented as mean ± SD. *Threshold values were estimated based on the maximum TSS criteria. The bimodality column displays the predictors that demonstrate a concave response curve. The name of variables was shortened as follows B = BIO.

| ID | Models | Threshold* | ROC-AUC | Sensitivity | Specificity | TSS | AICc | $\Delta$ AIC | $W_i$ | Bimodality |
| --- | --- | --- | --- | --- | --- | --- | --- | --- | --- | --- |
| 1 | B1 | 0.07 ± 0.01 | 0.87 ± 0.02 | 0.86 ± 0.02 | 0.73 ± 0.04 | 0.59 ± 0.03 | 2509.56 | 323.88 | 0.00 | |
| 2 | B12 | 0.09 ± 0.02 | 0.69 ± 0.03 | 0.76 ± 0.16 | 0.54 ± 0.18 | 0.30 ± 0.05 | 3164.57 | 978.89 | 0.00 | |
| 3 | I(B1<sup>2</sup>) | 0.06 ± 0.01 | 0.86 ± 0.02 | 0.86 ± 0.03 | 0.73 ± 0.03 | 0.58 ± 0.03 | 2599.64 | 413.96 | 0.00 | I(B1<sup>2</sup>) |
| 4 | I(B12<sup>2</sup>) | 0.10 ± 0.02 | 0.69 ± 0.03 | 0.76 ± 0.16 | 0.54 ± 0.18 | 0.30 ± 0.04 | 3191.21 | 1005.53 | 0.00 | |
| 5 | B1:B12 | 0.09 ± 0.00 | 0.65 ± 0.03 | 0.71 ± 0.09 | 0.58 ± 0.11 | 0.29 ± 0.04 | 3373.98 | 1188.30 | 0.00 | |
| 6 | B1 + B12 | 0.08 ± 0.01 | 0.90 ± 0.02 | 0.85 ± 0.03 | 0.82 ± 0.03 | 0.67 ± 0.04 | 2334.00 | 148.32 | 0.00 | |
| 7 | B1 + I(B1<sup>2</sup>) | 0.07 ± 0.02 | 0.87 ± 0.02 | 0.87 ± 0.03 | 0.73 ± 0.05 | 0.60 ± 0.03 | 2464.79 | 279.11 | 0.00 | |
| 8 | B1 + I(B12<sup>2</sup>) | 0.09 ± 0.01 | 0.90 ± 0.02 | 0.84 ± 0.03 | 0.83 ± 0.03 | 0.68 ± 0.04 | 2308.20 | 122.52 | 0.00 | |
| 9 | B1 + B1:B12 | 0.08 ± 0.01 | 0.90 ± 0.02 | 0.85 ± 0.03 | 0.82 ± 0.04 | 0.67 ± 0.04 | 2370.91 | 185.23 | 0.00 | |
| 10 | B12 + I(B1<sup>2</sup>) | 0.08 ± 0.00 | 0.90 ± 0.02 | 0.85 ± 0.04 | 0.82 ± 0.01 | 0.66 ± 0.05 | 2431.46 | 245.78 | 0.00 | I(B1<sup>2</sup>) |
| 11 | B12 + I(B12<sup>2</sup>) | 0.09 ± 0.03 | 0.69 ± 0.03 | 0.76 ± 0.16 | 0.54 ± 0.18 | 0.30 ± 0.05 | 3163.68 | 978.00 | 0.00 | I(B12<sup>2</sup>) |
| 12 | B12 + B1:B12 | 0.10 ± 0.02 | 0.89 ± 0.02 | 0.82 ± 0.03 | 0.82 ± 0.03 | 0.65 ± 0.06 | 2289.81 | 104.13 | 0.00 | |
| 13 | I(B1<sup>2</sup>) + I(B12<sup>2</sup>) | 0.08 ± 0.01 | 0.90 ± 0.02 | 0.84 ± 0.04 | 0.83 ± 0.03 | 0.67 ± 0.05 | 2411.86 | 226.18 | 0.00 | I(B1<sup>2</sup>) |
| 14 | I(B1<sup>2</sup>) + B1:B12 | 0.07 ± 0.01 | 0.90 ± 0.02 | 0.86 ± 0.04 | 0.80 ± 0.04 | 0.67 ± 0.04 | 2500.62 | 314.94 | 0.00 | I(B1<sup>2</sup>) |
| 15 | I(B12<sup>2</sup>) + B1:B12 | 0.11 ± 0.01 | 0.90 ± 0.02 | 0.84 ± 0.03 | 0.84 ± 0.03 | 0.68 ± 0.04 | 2282.01 | 96.33 | 0.00 | |
| 16 | B1 + B12 + I(B1<sup>2</sup>) | 0.09 ± 0.02 | 0.90 ± 0.02 | 0.86 ± 0.04 | 0.81 ± 0.04 | 0.68 ± 0.04 | 2257.71 | 72.03 | 0.00 | |
| 17 | B1 + B12 + I(B12<sup>2</sup>) | 0.09 ± 0.01 | 0.90 ± 0.02 | 0.85 ± 0.04 | 0.83 ± 0.03 | 0.68 ± 0.04 | 2295.67 | 109.99 | 0.00 | |
| 18 | B1 + B12 + B1:B12 | 0.10 ± 0.01 | 0.89 ± 0.02 | 0.82 ± 0.03 | 0.83 ± 0.03 | 0.65 ± 0.05 | 2281.90 | 96.22 | 0.00 | |
| 19 | B1 + I(B1<sup>2</sup>) + I(B12<sup>2</sup>) | 0.10 ± 0.02 | 0.90 ± 0.02 | 0.85 ± 0.05 | 0.83 ± 0.03 | 0.68 ± 0.04 | 2227.10 | 41.42 | 0.00 | |
| 20 | B1 + I(B1<sup>2</sup>) + B1:B12 | 0.09 ± 0.02 | 0.89 ± 0.01 | 0.86 ± 0.04 | 0.81 ± 0.03 | 0.67 ± 0.03 | 2287.32 | 101.64 | 0.00 | |
| 21 | B1 + I(B12<sup>2</sup>) + B1:B12 | 0.10 ± 0.01 | 0.90 ± 0.02 | 0.83 ± 0.04 | 0.85 ± 0.03 | 0.68 ± 0.04 | 2243.06 | 57.38 | 0.00 | |
| 22 | B12 + I(B1<sup>2</sup>) + I(B12<sup>2</sup>) | 0.08 ± 0.00 | 0.90 ± 0.02 | 0.84 ± 0.04 | 0.84 ± 0.03 | 0.68 ± 0.04 | 2407.17 | 221.49 | 0.00 | I(B1<sup>2</sup>) |
| 23 | B12 + I(B1<sup>2</sup>) + B1:B12 | 0.10 ± 0.02 | 0.89 ± 0.02 | 0.83 ± 0.03 | 0.82 ± 0.03 | 0.65 ± 0.06 | 2291.80 | 106.12 | 0.00 | I(B1<sup>2</sup>) |
| 24 | B12 + I(B12<sup>2</sup>) + B1:B12 | 0.10 ± 0.00 | 0.89 ± 0.02 | 0.84 ± 0.04 | 0.82 ± 0.01 | 0.66 ± 0.05 | 2241.64 | 55.96 | 0.00 | |
| 25 | I(B1<sup>2</sup>) + I(B12<sup>2</sup>) + B1:B12 | 0.11 ± 0.01 | 0.90 ± 0.02 | 0.83 ± 0.03 | 0.84 ± 0.03 | 0.68 ± 0.04 | 2268.56 | 82.88 | 0.00 | I(B1<sup>2</sup>) |
| 26 | B1 + B12 + I(B1<sup>2</sup>) + I(B12<sup>2</sup>) | 0.11 ± 0.02 | 0.90 ± 0.02 | 0.85 ± 0.04 | 0.83 ± 0.03 | 0.68 ± 0.04 | 2212.31 | 26.63 | 0.00 | |
| 27 | B1 + B12 + I(B1<sup>2</sup>) + B1:B12 | 0.09 ± 0.01 | 0.90 ± 0.02 | 0.86 ± 0.02 | 0.80 ± 0.02 | 0.67 ± 0.04 | 2226.49 | 40.81 | 0.00 | |
| 28 | B1 + B12 + I(B12<sup>2</sup>) + B1:B12 | 0.10 ± 0.01 | 0.90 ± 0.02 | 0.84 ± 0.04 | 0.82 ± 0.02 | 0.66 ± 0.05 | 2237.05 | 51.37 | 0.00 | |
| 29 | **B1 + I(B1<sup>2</sup>) + I(B12<sup>2</sup>) + B1:B12** | **0.10 ± 0.02** | **0.90 ± 0.02** | **0.86 ± 0.04** | **0.82 ± 0.03** | **0.68 ± 0.04** | **2186.70** | **01.02** | **0.38** |  |
| 30 | B12 + I(B1<sup>2</sup>) + I(B12<sup>2</sup>) + B1:B12 | 0.10 ± 0.01 | 0.89 ± 0.02 | 0.84 ± 0.05 | 0.82 ± 0.02 | 0.66 ± 0.05 | 2243.56 | 57.88 | 0.00 | |
| 31 | **B1 + B12 + I(B1<sup>2</sup>) + I(B12<sup>2</sup>) + B1:B12** | **0.10 ± 0.02** | **0.90 ± 0.02** | **0.86 ± 0.04** | **0.83 ± 0.02** | **0.68 ± 0.04** | **2185.68** | **0.00** | **0.62** |  |

Response curves for the two climatic variables presented a well-defined Gaussian shape (Fig. 1). The peak probability for BIO-1 occurs at around 20°C, whereas for BIO-12, the maximum probability was observed for values below 500 mm. Extrapolation of these curves to values outside calibration ranges showed decreasing probabilities, indicating safe model extrapolations rather than perpetual increments of probability towards extreme environmental conditions.

**Figure 1.**
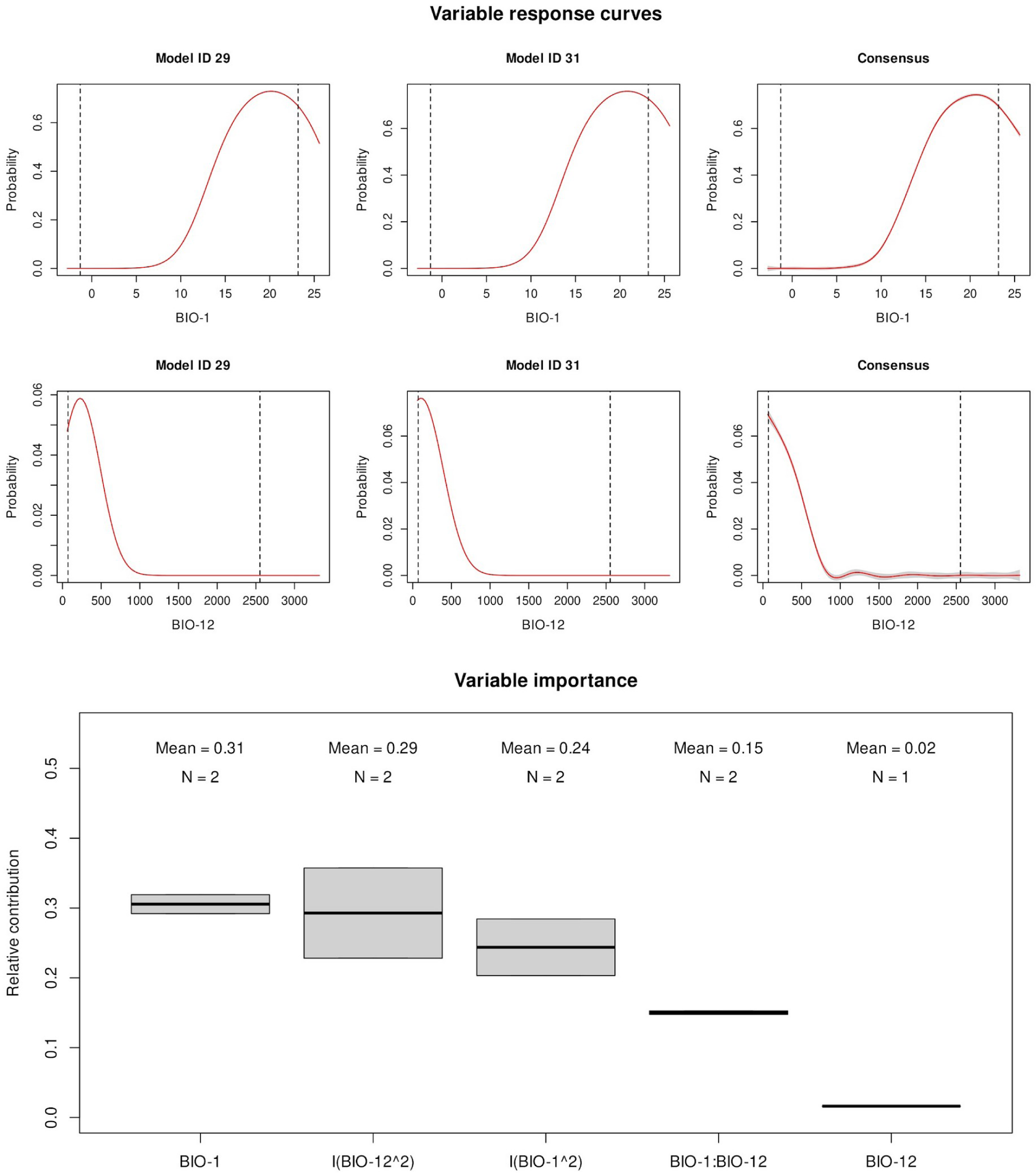
Bioclimatic variable response and importance across models. The upper half illustrates the response curves of the species to variables BIO-1 (annual mean temperature °C) and BIO-12 (Annual precipitation mm) across the two best models (ID 29 and ID 31), and the vertical dashed lines mark environmental limits of the calibration data. The lower half of the figure shows a boxplot detailing the distribution of variable importance values among selected models and highlighting the variability and median importance of predictors. The importance of variables, measured by explained deviance, incorporates linear, quadratic terms (denoted as I(var^2)) and two-way interaction. The y-axis represents the importance values, with boxplots delineating the interquartile range and median value. The frequency of predictor inclusion across models is also noted, providing insight into their relative significance.

Variable importance analysis highlighted the linear term of BIO-1 and the quadratic terms of both variables as the most significant predictors (Fig. 1). The interaction between variables, along with the linear component of BIO-12, were found to be less relevant. Despite their minor contribution, models incorporating these predictors were selected based on their superior goodness of fit compared to models excluding them.

The geographic predictions of the two selected models showed consistent patterns, with higher probability values in the southwestern parts of the United States (Fig. 3). However, some discrepancies were noted in Florida, where one model estimates higher probability values than the other (Fig. 2). Independent data validation confirmed the comparable performance of all projected models, although with slightly different threshold values estimated for each model (Tables 3 and S1).

**Table 3.** Evaluation of the two selected models and the consensus using an independent data set with presence and absence records. The classification capacities of the final prediction were calculated using the confusion matrix based on three threshold criteria.

| Model | Threshold criteria | Threshold | ROC-AUC | False positive rate | Accuracy | Sensitivity | Specificity | TSS |
| --- | --- | --- | --- | --- | --- | --- | --- | --- |
| ID 29 | ESS | 0.137 | 0.957 | 0.090 | 0.910 | 0.909 | 0.910 | 0.819 |
|  | maxTSS | 0.117 | 0.957 | 0.101 | 0.910 | 1.000 | 0.899 | 0.899 |
|  | SEN90 | 0.131 | 0.957 | 0.101 | 0.900 | 0.909 | 0.899 | 0.808 |
| ID 31 | ESS | 0.158 | 0.957 | 0.090 | 0.910 | 0.909 | 0.910 | 0.819 |
|  | maxTSS | 0.135 | 0.957 | 0.101 | 0.910 | 1.000 | 0.899 | 0.899 |
|  | SEN90 | 0.144 | 0.957 | 0.101 | 0.900 | 0.909 | 0.899 | 0.808 |
| Consensus (weighted average) | ESS | 0.150 | 0.957 | 0.090 | 0.910 | 0.909 | 0.910 | 0.819 |
|  | maxTSS | 0.127 | 0.957 | 0.101 | 0.910 | 1.000 | 0.899 | 0.899 |
|  | SEN90 | 0.139 | 0.957 | 0.101 | 0.900 | 0.909 | 0.899 | 0.808 |

**Figure 2.**
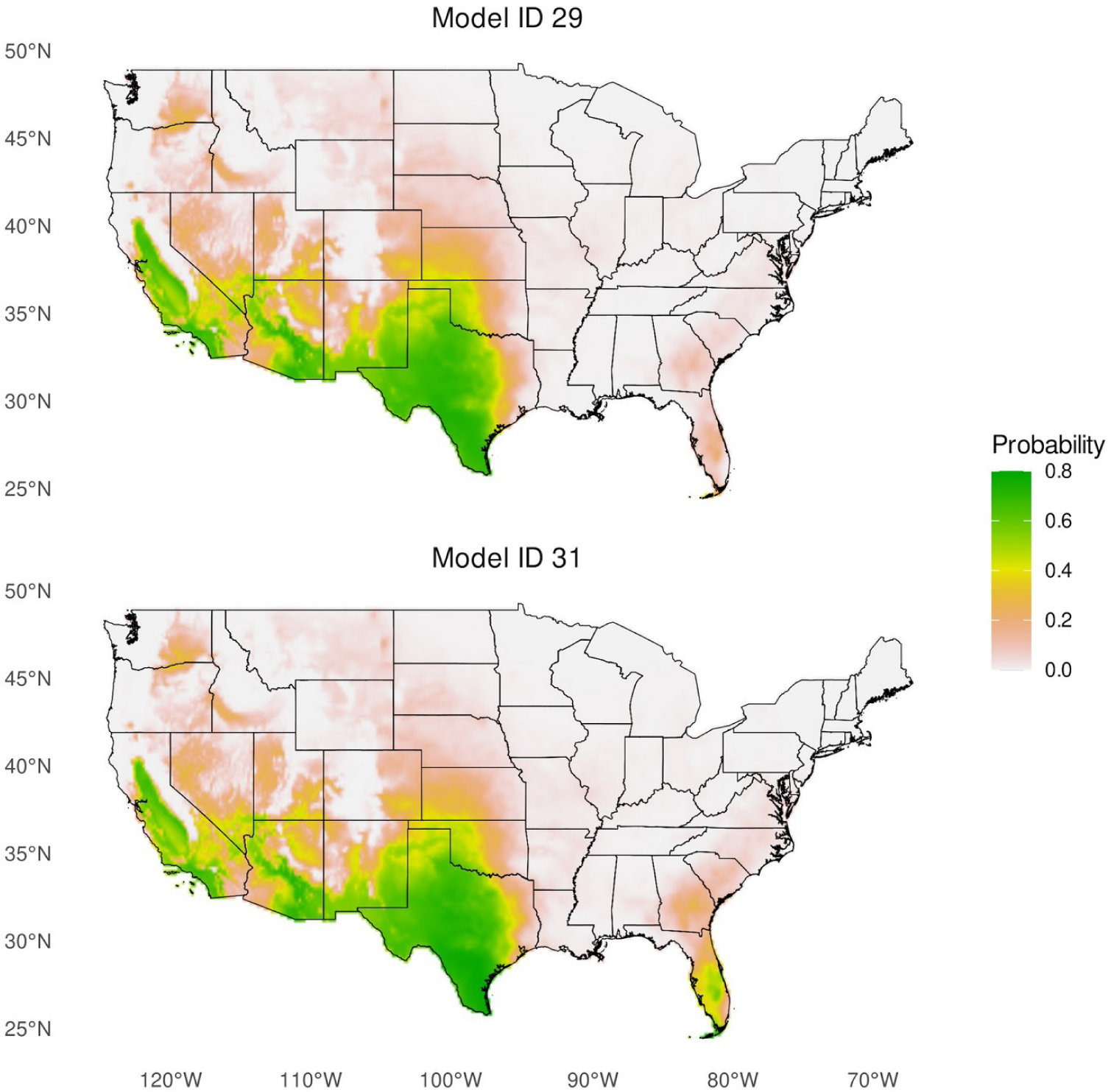
Geographic depiction of estimated probability of occurrence for the pathogen virtual species deriving from the two final selected models. Maps are shown at a spatial resolution of 10’ (∼20 km at the Equator).

**Figure 3.**
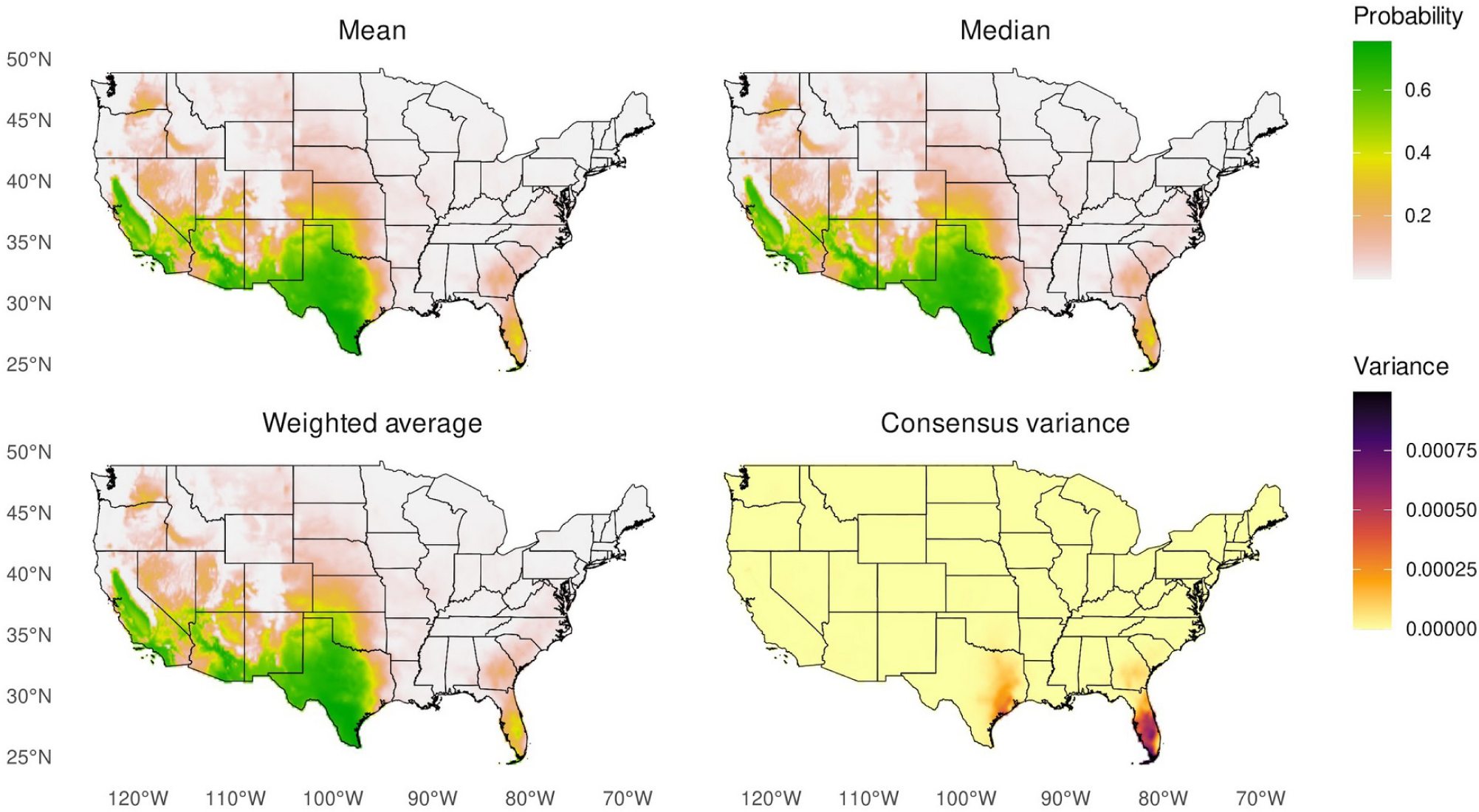
Consensus geographic projection of probability of occurrence for the pathogen virtual species. The figure displays the probability of occurrence derived from the two final selected models using averaging metrics: mean, median, and weighted average based on Akaike weights. In this example, since the averaging is calculated from two models, the median coincides with the mean. The figure displays the variance among the three consensus projections. The maps are shown at a spatial resolution of 10‘ (∼20 km at the Equator).

## Discussion

Understanding how species are distributed in different environments and predicting how species will respond to changes is crucial in research in ecology. Ecological niche modeling helps in this task, and this contribution introduces enmpa, an R package that facilitates calibration of ecological niche models via GLM. The package integrates a set of complex methodological developments in the ENM field, using logistic GLMs to estimate the probability of a species occurring in a particular environment (Austin 2002, 2007; Ward et al. 2009). The tools developed in this package are particularly interesting for studies that involve data derived from detection/non-detection sampling protocols, such as pathogen test results, detections of species on controlled-protocol surveys, etc.

One of the most notable features of enmpa is that it allows users to explore a wide range of predictor combinations in a GLM framework to find the set of combinations that better fit the data and explain the phenomenon of study. Focusing on exploring different predictor features, including linear, quadratic, and two-way interaction responses, enabling a detailed analysis of the entire parameter space (Cobos et al. 2019b).

Apart from the functionalities corresponding to the main steps in ENM, enmpa implements two novel methods that allow users to select variables based on niche signal detection (Cobos and Peterson 2022) and filter those models with response shapes that do not align with ecological theory (Austin 2002, 2007; Peterson et al. 2011; Merow et al. 2014). The first method helps users to discard potentially non-significant variables before modeling, which prevents over-parameterization of models and makes the calibration step easier by avoiding exploring irrelevant predictor combinations that do not form part of the species’ niche (Cobos and Peterson 2022). Although forward, backward, or stepwise selection processes have been commonly used methods for selecting predictors in modeling (Efroymson 1960), they have been criticized for misapplication of a single-step statistical test in a multi-step procedure (Harrell 2001; Flom and Cassell 2007; Smith 2018). This problem may lead to the selection of nuisance variables or models that perform worse with independent data than in calibration (Smith 2018).

The second method seeks to stay in line with the standard of niche theory, in which species’ fitness responds to environmental conditions with a unimodal response. Extreme environmental values lead to low fitness, while intermediate environmental values are optimal for the species (Jiménez-Valverde et al. 2011; Escobar 2020). However, the fitting of quadratic terms can be limited if insufficient sampling information is available and one extreme of the curve is not captured, which can lead to estimation of odd response shapes. Although bimodal responses of species to variables may have interesting and meaningful interpretations (e.g., indicating that a niche is incompletely represented by the data), they need to be evaluated in detail for particular species. Therefore, excluding quadratic predictors with bimodal behavior is crucial in most cases to avoid misleading conclusions.

The example of a virtual pathogen species demonstrates the usefulness and effectiveness of enmpa. The meticulous steps involved in model calibration, selection, and evaluation, in combination with the consideration of response curves and variable contributions, collectively contributed to a refined understanding of the ecological niche of the virtual species. For example, the results suggest that the virtual pathogen would thrive in warmer climates with lower rainfall. The example also highlights the practicality and accuracy of enmpa in modeling species’ niches, when records derived from sampling protocols of detection and non-detection are available.

## Acknowledgments

We would like to thank the KUENM working group for valuable discussions. LFAG would like to thank the Biodiversity Institute, University of Kansas, for hosting him during the development of this project. LFAG thanks the support of Juan A. Navas-Cortés and Blanca B. Landa during the development of this work.

## Supplementary materials

**Figure S1.**
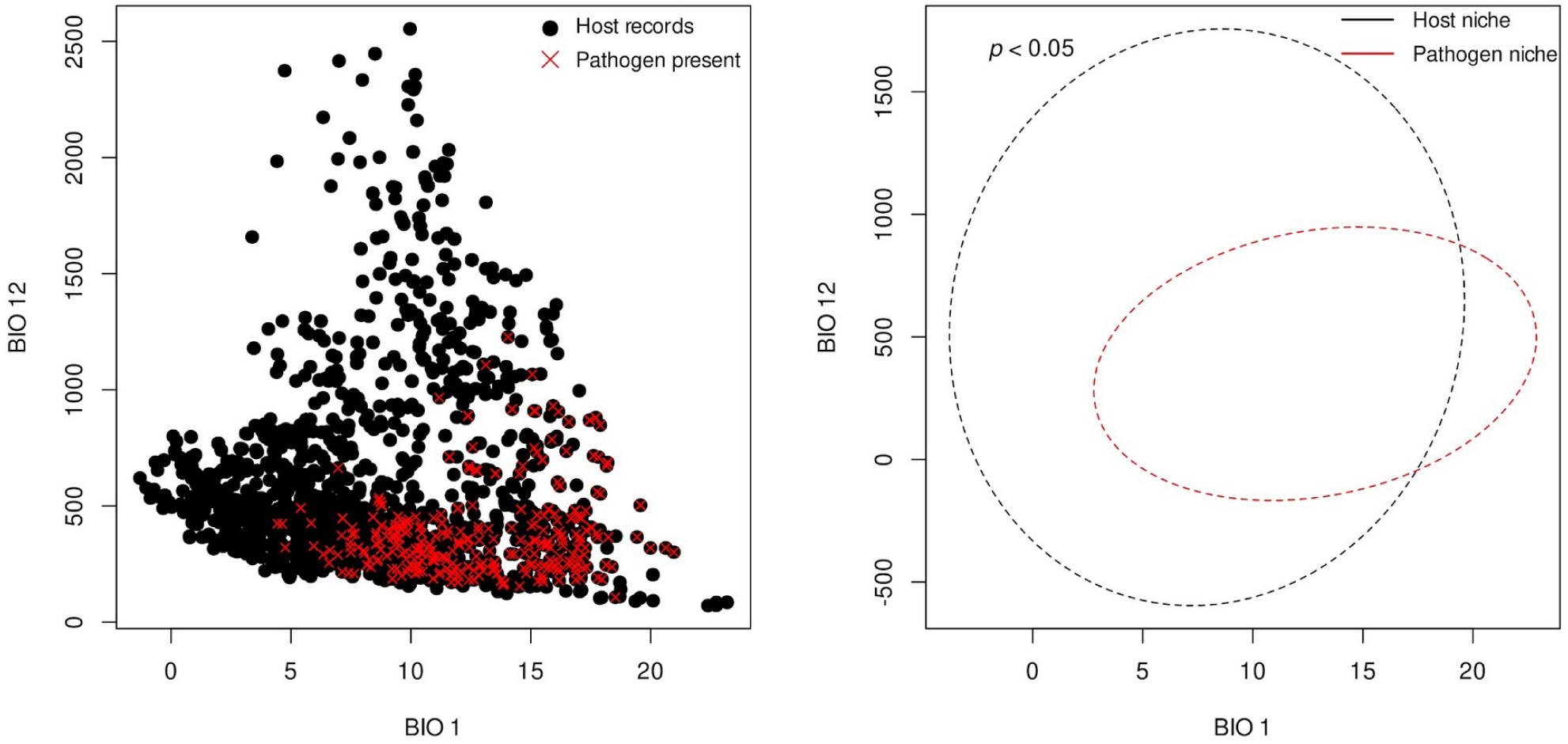
Results from niche comparison using PERMANOVA analysis. The figure on the left represents positive and negative records in the environmental space for BIO 1 and BIO 2. On the right, ellipsoids are derived from the data to explore and visualize the position and spread of host and pathogen niches. The *p*-value represents the statistical significance of the PERMANOVA test.

**Figure S2:**
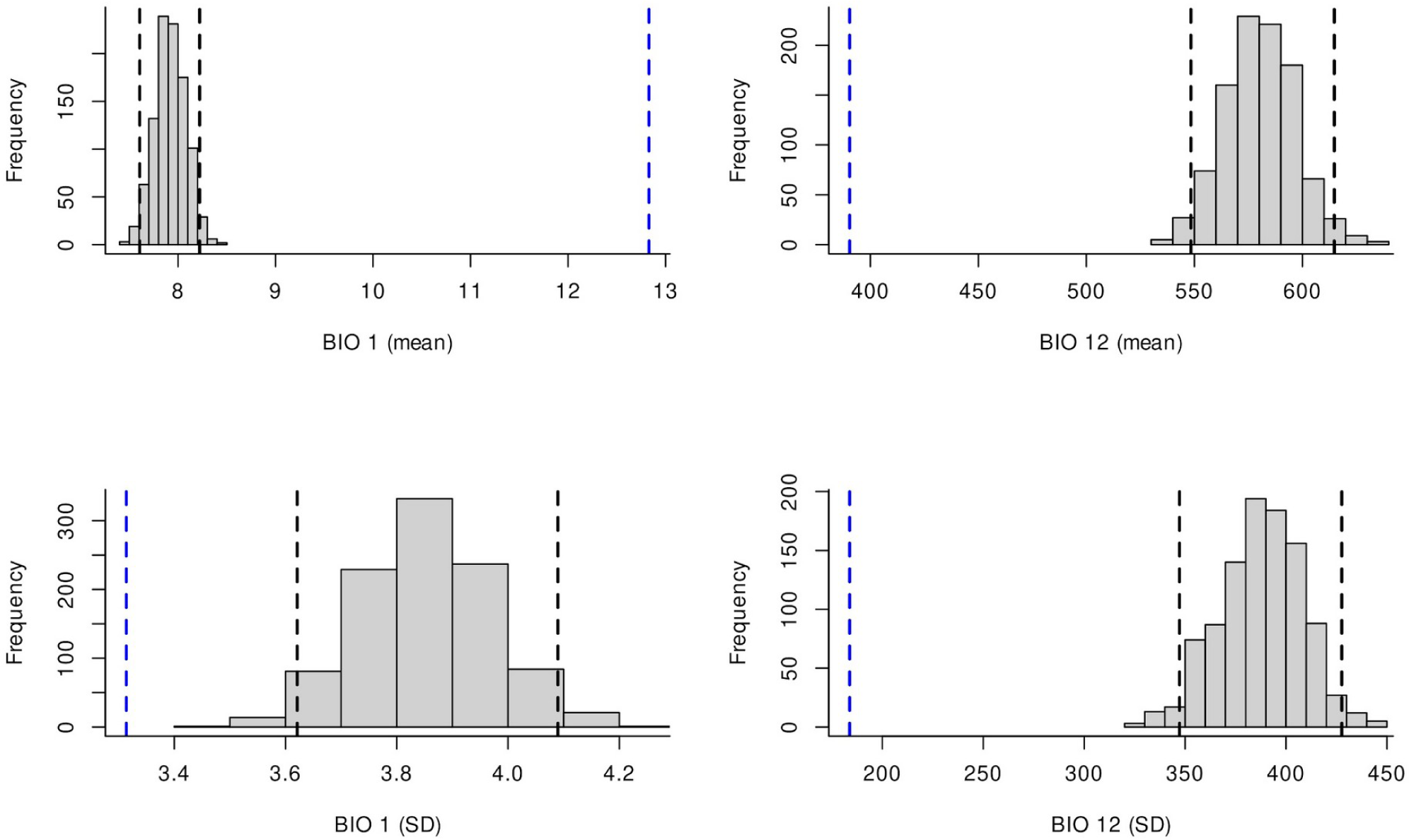
Visualization of the results obtained from the univariate non-parametric test for detecting signals of the virtual pathogen’s niche. The top-left and top-right panels depict the mean distribution of the niche position of the species about the null distribution for the BIO 1 and BIO 12 variables. Bottom-left and bottom-right panels depict the range of environmental conditions of the niche about the null distribution, as represented by the standard deviation (SD). The vertical dotted blue lines signify the observed value associated with presences of the pathogen. The vertical dotted gray lines represent the lower and upper 95% confidence limits of the null distribution. The barplot histogram represents the null distribution.

**Table S1.** Evaluation of the two selected models and the consensus using an independent data set with presence and absence records. Using presences-only.

| Models | Threshold criteria | Threshold | Omission error | Mean AUC ratio at 5% | P-value pROC |
| --- | --- | --- | --- | --- | --- |
| ID 29 | ESS | 0.137 | 0.091 | 1.648 | 0.000 |
|  | maxTSS | 0.117 | 0.000 | 1.648 | 0.000 |
|  | SEN90 | 0.131 | 0.091 | 1.648 | 0.000 |
| ID 31 | ESS | 0.158 | 0.091 | 1.631 | 0.000 |
|  | maxTSS | 0.135 | 0.000 | 1.630 | 0.000 |
|  | SEN90 | 0.144 | 0.091 | 1.636 | 0.000 |
| Consensus (weighted average) | ESS | 0.150 | 0.091 | 1.635 | 0.000 |
|  | maxTSS | 0.127 | 0.000 | 1.640 | 0.000 |
|  | SEN90 | 0.139 | 0.091 | 1.640 | 0.000 |

